# Molecular phenotyping using networks, diffusion, and topology: soft tissue sarcoma

**DOI:** 10.1101/328054

**Authors:** James C Mathews, Maryam Pouryahya, Caroline Moosmüller, Ioannis Kevrekidis, Joseph O Deasy, Allen Tannenbaum

## Abstract

Many biological datasets are high-dimensional yet manifest an underlying order. In this paper, we describe an unsupervised data analysis methodology that operates in the setting of a multivariate dataset and a network which expresses influence between the variables of the given set. The technique involves network geometry employing the Wasserstein distance, global spectral analysis in the form of diffusion maps, and topological data analysis using the Mapper algorithm. The prototypical application is to gene expression profiles obtained from RNA-Seq experiments on a collection of tissue samples, considering only genes whose protein products participate in a known pathway or network of interest. Employing the technique, we discern several coherent states or signatures displayed by the gene expression profiles of the sarcomas in the Cancer Genome Atlas along the p53 signaling network. The signatures substantially recover the leiomyosarcoma, dedifferentiated liposarcoma (DDLPS), and synovial sarcoma histological subtype diagnoses, but they also include a new signature defined by simultaneous activation and inactivation of about a dozen genes, including activation of fibrinolysis inhibitor SERPINE1/PAI and inactivation of p53-family tumor suppressor gene P73 along with cyclin dependent kinase inhibitor 2A CDKN2A/P14ARF.

## 1 Introduction and Background

Modern biological investigations often result in dense, high-dimensional datasets describing genes, proteins, mutations, or other variables. A near universal problem arises as the dimensionality of the data grows: How can the data be investigated in a relatively unbiased manner, to expose underlying clusters and relational structure? To date, there is a lack of robust techniques for exposing the structure of biological data in an unbiased, agnostic fashion. Biological relevance is maintained in this work by considering known pathways as an underlying guide. The technique we describe involves network geometry via the Wasserstein distance ([Rachev and Rüschendorf, 1998], [Rubner et al., 2000]), global spectral analysis in the form of diffusion maps ([Coifman and Lafon, 2006]), and topological data analysis using the Mapper algorithm ([Nicolau et al., 2011]). The prototypical application is to gene expression profiles obtained from RNA-Seq experiments on a collection of tissue samples, considering only genes whose protein products participate in a known pathway or network of interest. Using the technique, we discern several coherent states or signatures displayed by the gene expression profiles of the sarcomas in the Cancer Genome Atlas along the p53 signaling network. The signatures substantially recover the leiomyosarcoma, dedifferentiated liposarcoma (DDLPS), and synovial sarcoma histological subtype diagnoses, but they also include a new signature defined by simultaneous activation and inactivation of about a dozen genes (including activation of fibrinolysis inhibitor SERPINE1/PAI and inactivation of p53-family tumor suppressor gene P73 and cyclin dependent kinase inhibitor 2A CDKN2A/P14ARF).

The mechanisms that intervene between DNA sequence genotype and overall cell phenotype are clearly very complex, including the presence of transcription factors, the chemistry of the cell microenvironment, and epigenetic factors like methylation, acetylation, or ubiquitination. Leaving aside the elucidation of these mechanisms *per se*, one first needs techniques for making qualitative distinctions between the states that are the most immediate and most accessible effects of these mechanisms, typically understood in terms of experimentally determined or inferred gene expression quantifications (ideally, counts or concentration levels of RNA transcripts or proteins). Such distinctions may be called “molecular phenotypes”. Molecular phenotypes are apparent in contexts where the abundance of a small number of gene products is known to be strongly correlated with particular outcomes of interest, e.g., in breast cancer studies where elevated presence of the estrogen receptor protein (ER), the progesterone receptor protein (PR), or the human epidermal growth factor receptor 2 protein (HER2/ERBB2) are markers for amenability to certain hormone therapies. However, in general such a marker protein may not exist, and one must instead measure, somehow, the cumulative effect of a comparatively large number of gene products.

Methods falling under the heading of Genome Wide Association Studies (GWAS, typically concerning mutational profiles) or Gene Set Enrichment Analysis (GSEA, typically concerning gene expression profiles) take into account data concerning a large number of genes to ascertain statistical significance with respect to given known outcomes or endpoints such as disease states. In general, they do not attempt to discern coherent states in gene expression quantification profiles in an “unsupervised” manner. That is, these methods do not ascertain existing apparent molecular phenotypes, but rather impose or design molecular phenotypes specifically to serve as predictors for variables of ultimate interest like prognosis. Though designed predictors are indeed indispensable for specific urgent tasks, for long-term reusability and overall scientific value one seeks predictors grounded in observations made directly from data in light of known mechanisms.

In [Seemann et al., 2012], the authors employ degree zero persistent homology towards this end, leveraging the robustness of Topological Data Analysis (TDA) techniques for unsupervised clustering. One could also use various established unsupervised clustering algorithms such as hierarchical clustering or *k*- mean optimization methods, optionally preceded by dimensional reduction techniques like Principal Component Analysis (PCA), *t*-distributed Stochastic Neighbor Embedding (t-SNE), or Multi-Dimensional Scaling (MDS). Note, however, that hierarchical clustering has the drawback that the output of the algorithm strongly under determines the usual heat map visual representation. Every branch of the hierarchy tree creates an ambiguity in the order in which the samples are displayed.

From the topological point of view, however, any method within the clustering paradigm is order zero in the sense that it summarizes a dataset with discrete categorization in terms of a finite set of disjoint categories, a “space” of dimension zero. In [Lockwood and Krishnamoorthy, 2014], the authors advocate “higher order” methods, e.g. degree one persistent homology, extracting one-dimensional features in the space charted by the data points, roughly in order to take account of the relations between categories and not just the categories themselves. One major difficulty with this approach is that homology classes are defined by *cycles*; topological features which are not cycles, such as *relative cycles* or branches, are not detectable with existing tools (see [Munkres, 1975] for general background on topology, or [Edelsbrunner and Harer, 2010] for background on TDA methods). A second major difficulty is that homology classes do not have canonical representative cycles. This means that in theory an almost arbitrary subset of the points of a point cloud can appear along the path of a cycle belonging to an observed persistent 1-homology class. In other words, while persistent homologies are certainly evidence of important dataset-specific global features, there is an unsolved problem of interpretability of such features.

The authors [Camara et al., 2016] calculate persistent 1-homologies in evolutionary/phylogenetic data, surmounting both of these difficulties simultaneously by interpreting the presence of non-trivial cycles (closed loops), and not the internal structure of their representative cycles *per se*, as an indication of the presence of genetic recombination events.

We largely follow [Nicolau et al., 2011] in that we use the Mapper algorithm to map our point clouds onto “summary” spaces of dimension one, graphs or networks. This algorithm can be regarded as a discrete version of the Morse-theoretic analysis of a smooth manifold with respect to a height function (called the filter function). Nicolau *et al.* heavily de-sparsify the point clouds, in order to avoid the normal pre-processing step of dimensional reduction (virtually always required for biological datasets), and employ a carefully designed deviation-from-normal filter function in accordance with what they call the Progression Analysis of Disease paradigm. We take a slightly different tack: First, we perform a biologically-motivated intermediate-scale dimensional reduction by considering only those genes participating in well-known pathways (we use the Kyoto Encyclopedia of Genes and Genomes). Next, we perform a replacement of the ordinary Euclidean distance metric between gene expression profiles with alternative metrics, especially a version of the Wasserstein 1-metric which takes account of curated knowledge of the network structure linking the genes (coordinates). Then we perform the further dimensional reduction and analysis technique of diffusion maps [Coifman and Lafon, 2006] to regularize the point cloud with respect to intrinsic or characteristic global geometry. We have found that this process results in datasets with favorable properties for the application of Mapper and interpretation of its resultant graph summaries.

## 2 Methods

We take as our primary input a gene expression quantification sample set, as a point cloud *S* ⊂ ℝ^*>N*^, an influence, regulation, or pathway network *G*_*g*_ relating the *N* genes which label the coordinates. Optionally, we include an additional control dataset *C* ⊂ ℝ^*>N*^, or a function *f*: *S* → ℝ with the interpretation as an experimentally-determined “degree of progression” with respect to some process (e.g., a disease process).

The output is a list of coherent states or molecular phenotypes, characterized by activated, inactivated, and equivocally-activated genes. We now enumerate the steps of our pipeline. Details will be given in subsequent sections.

1. Normalizing the values of *S*
2. Restricting (projecting) *S* to the genes appearing in the gene network *G*_*g*_
3. Calculation of a network-based distance metric between samples
4. Evaluation of a diffusion map
5. The Mapper algorithm
6. State graph extraction and processing
7. Plotting heat maps and discerning coherent states

### 2.1 Normalizing the values of *S*

We must ensure that the values of *S* represent gene expression quantification, for example, FPKM (Fragments Per Kilobase Million) or TPM (Transcripts Per Kilobase) values as a result of a high-throughput sequencing pipeline. These values correspond roughly to the concentration of RNA transcripts in the tissue samples, typically across many cells for each sample (bulk sequencing) though sometimes for single cells. With no additional information we may reasonably assume that transcript levels approximately correspond to protein product concentrations.

Optionally, for each of the *N* genes, we replace the values of *S* in the corresponding coordinate by a truncated translated *z*-score:

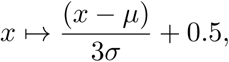

where *µ* and *σ* are the mean and standard deviation of the values for this coordinate and *x* is a typical value of this coordinate. The resulting values will be substantially normalized to lie on a scale from 0 to 1 with mean 0.5. Threshold any negative values to 0.

### 2.2 Restricting (projecting) *S* to the genes appearing in the gene network *G*_*g*_

We select a network *G*_*g*_whose nodes correspond to genes whose presence or absence constitutes participation in a coordinated function or process of interest, and whose edges represent the coordinating relationships. Omit the expression values for genes not participating in *G*_*g*_. The networks we consider are the KEGG (Kyoto Encyclopedia of Genes and Genomes) pathways concerning cell cycle regulation, senescence, proliferation, apoptosis, and p53 signaling.

### 2.3 Calculation of a network-based distance metric between samples

For each sample *s* ∈ *S*, define a probability distribution *p*_*s*_on the set of nodes of *G*_*g*_by interpreting the values of *s* (divided by their sum) as a probability density function. Alternatively, use a distribution on the nodes which is the invariant measure for a Markov chain stochastic process inferred from the values of the sample *s*, in the manner of [Chen et al., 2017]. In case *G*_*g*_consists of several connected components, define separate distributions for each component. For each component *c*, calculate the Wasserstein 1-metric *d*_*c*_(*s, s*′) between each pair *p*_*s*_ and *p*, _*s*_ *′* also known as the Earth-Mover′s Distance, with respect to the pathlength metric on *c* (weighted by the reciprocal of strength-related edge weights, if present). The Earth Mover′s Distance between two mass distributions on a common metric space is defined as the infimum of the total mass-weighted displacement over displacement functions from the space to itself which map the first mass distribution onto the second. Classically it is only defined if the total masses of both distributions are equal; see [Rachev and Rüschendorf, 1998] and the references therein. We use the “direct sum” formula to amalgamate these distances across components *c* into a single distance for each pair of samples (*s, s*′):

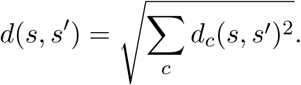

The Wasserstein 1-metric employed in this way is perhaps the simplest alternative to the standard Euclidean metric for which a network or pathway structure relating the coordinates is in some way taken into account. The principal benefit of this metric is that it greatly increases the distance between two samples in comparison with the Euclidean metric in case the main activity of one sample takes places in an area of the network very far from the area of main activity of the other sample. One conceivable disadvantage is that isolated changes to a given sample, say in the expression of a single gene, can have an outsized effect on the Wasserstein 1-distance of the displacement. We also caution that since the KEGG database networks are enriched with nodes for various compounds and macromolecules in addition to protein gene products, an analysis which considers only gene expressions will not take advantage of the full KEGG pathways and may have some misleading consequences. To compute the Wasserstein distance, we used the Hungarian algorithm of [Rubner et al., 2000].

**Figure 1:**
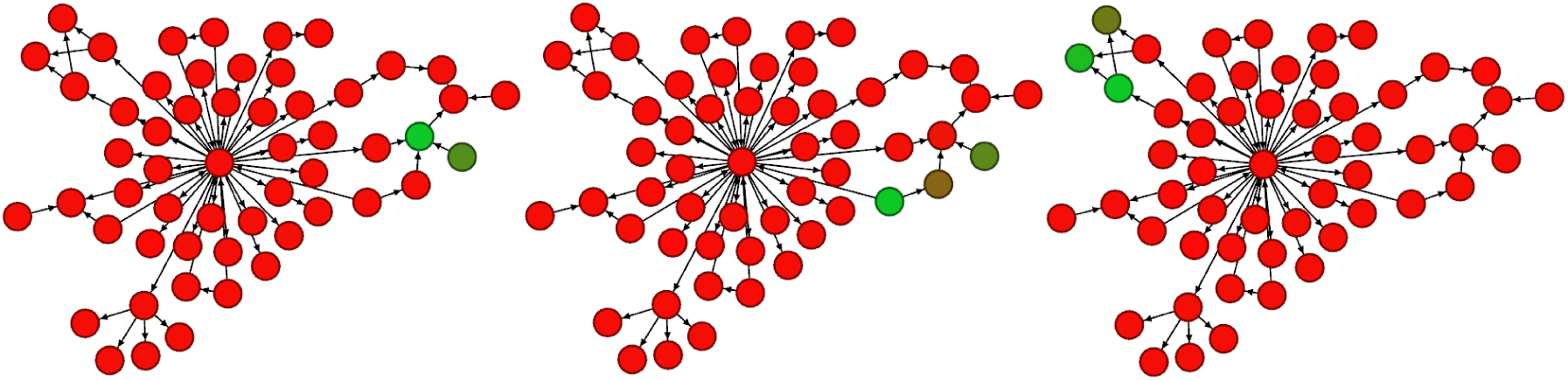
Left to right, distributions *s*_1_, *s*_2_, *s*_3_, for illustration. Red represents values close to zero, and green more positive. The Wasserstein 1-distance *d*(*s*_1_, *s*_2_) = 1.10 is much less than *d*(*s*_2_, *s*_3_) = 3.08, while the corresponding Euclidean distances are approximately equal to each other.

### 2.4 Evaluation of a diffusion map

In order to reduce the dimension and complexity of the dataset *S*, while preserving key information for subsequent analysis, we apply diffusion maps [Coifman et al., 2005, Coifman and Lafon, 2006]. This manifold learning technique provides a global parametrization of a low-dimensional, possibly nonlinear manifold on which the high-dimensional data is assumed to lie. Such an embedding is obtained by spectral properties (eigenvalues and eigenvectors) of the graph Laplacian on a certain weighted graph with nodes *S*. Its eigenvectors can be used as a coordinate system on the dataset *S*, which is justified by the fact that they approximate the eigenvectors of the Laplace-Beltrami operator of the underlying manifold [Coifman and Lafon, 2006, Jones et al., 2008]. See also some recent work [Rajendran 2016] for employing diffusion map techniques for data whose “points” are weighted graphs.

For data samples *s*_*i*_, *s*_*j*_ ϵ *S* we define a connectivity matrix *W* using a Gaussian kernel:

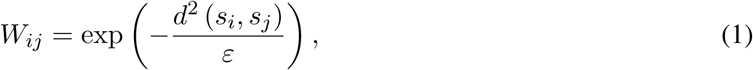

where *d* is the Wasserstein 1-metric as defined in [Rachev and Rüschendorf, 1998] and *ε* is the kernel scale parameter. The kernel is intended to capture the features of the underlying dataset and it is therefore reasonable to choose the metric *d* and the scale parameter *ε* based on the application. The parameter *ε* defines a local connectivity scale, in the following sense: If *s*_*j*_ is in the *ε*-ball around *s*_*j*_ the kernel induces high weight between *s*_*i*_ and *s*_*j*_. Otherwise the weights are negligible. We can choose *ε* to be almost any value between the minimum and maximum among the pairwise squared distances (*d*(*s*_*j*_ *s*_*j*_))^2^.

Define an adapted kernel

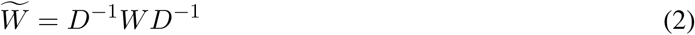

With D the diagonal matrix 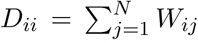, *N* = #*S* We use the adapted kernel (2) instead of (1), corresponding to the choice *α* = 1 in the family of kernel normalizations presented in [Coifman and Lafon, 2006, Nadler et al., 2006], to recover the Riemannian geometry of the underlying data independently of the data sampling.

We build a weighted graph with node set *S* and weights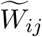 of the edge connecting *s*_*i*_ to *s*_*j*_. Now apply weighted graph Laplacian normalization to 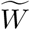

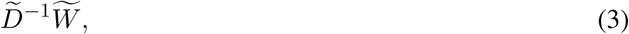

With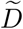 the diagonal matrix 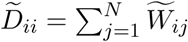 The associated graph Laplacian is given by

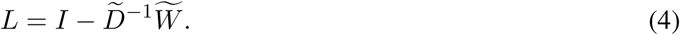

We compute the eigenvalues 1 = *λ*_1_ ≥ |*λ*_2_| ≥…≥ |*λ*_*N*_*|* and eigenvectors *ø*_1_, …, *ø*_*N*_ of *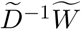*. These eigenvectors provide an embedding of the data into a space of dimension *M* < *N*:

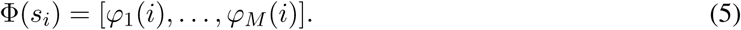

**Figure 2:**
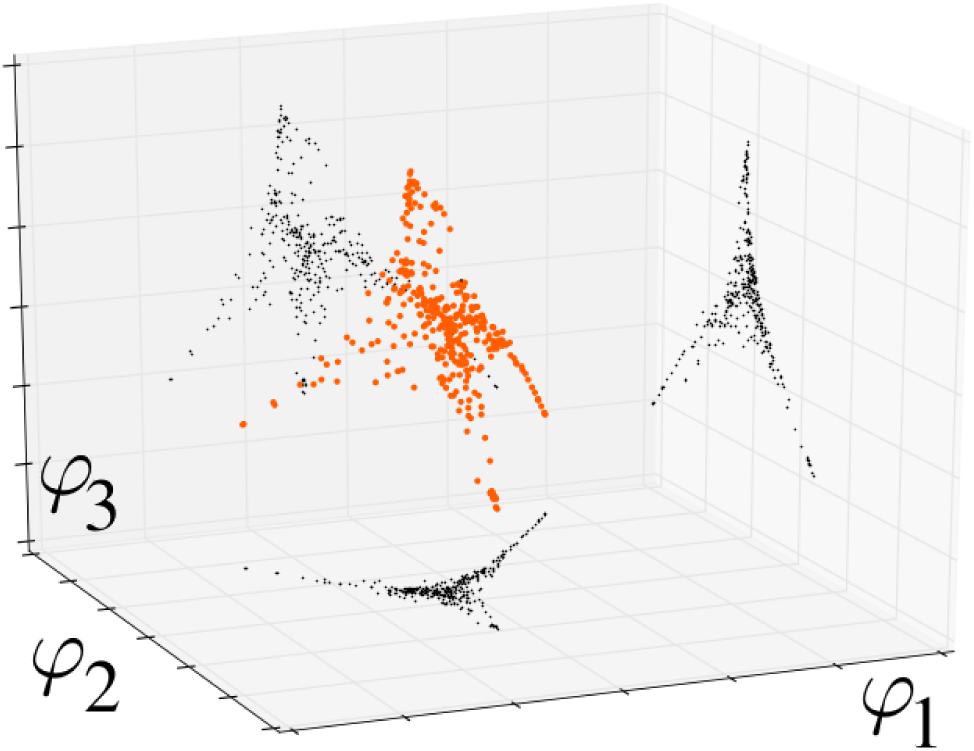
The diffusion re-mapped images of the gene expressions of 355 adipose visceral omentum tissue samples from the GTEx database. The first three eigenfunctions of the diffusion operator are used, to make a three-dimensional plot.

We reiterate that rather than using the Euclidean distance between samples *s*_*i*_ and *s*_*j*_, we select the “more informed” 1-Wasserstein network metric. Unlike dimensional reduction techniques like Principal Component Analysis or Local Linear Embedding, but in common with *t*-SNE or MDS, the technique of diffusion maps can function on arbitrary intrinsic-metric representations of the data of the point cloud and does not require this point cloud to be presented in some Euclidean space. We prefer diffusion maps over t-SNE or MDS because as far as we know the latter are not guaranteed to recover the intrinsic manifold degrees of freedom of the original dataset, while diffusion maps are so guaranteed in principle. In practice biological datasets of present interest seem to represent processes of sufficient complexity that precise quantitative accounting for the relationships between all of the variables is rarely proposed, so such theoretical considerations are arguably premature.

We remark that dimensional reduction of gene expression data via diffusion maps is also suggested e.g. in [Xu et al., 2010], where the authors combine diffusion maps with a neural network clustering method to differentiate between different types of small round blue-cell tumors.

### 2.5 The Mapper algorithm

This algorithm results in a simplicial complex, in some sense modeling the mesoscopic-scale topology of the support space for the collection of samples or states *S*. It works by (1) dividing the point cloud into overlapping “slices” by binning the values of a chosen filter function *f*: *S →* ℝ into overlapping bins, (2) clustering the points of each slice (e.g. with single-linkage clustering), and (3) linking pairs (or tuples) of clusters by edges (or higher-dimensional simplices) depending on the amount of overlap between clusters.

A reasonable choice for the filter function is a deviation function devised in comparison with a control dataset *C*, roughly as in [Nicolau et al., 2011], e.g. the Mahalanobis distance function adapted to *C* in case the size of *C* is sufficiently large in comparison to the dimension *M*. We often use the general-purpose network centrality measure available in Daniel Müllner′s Python Mapper implementation^1^.

The algorithm requires the choice of certain resolution or scale parameters: the number *n*_*f*_ of filter-level-set bins and a threshold *t* for the single-linkage clustering algorithm applied to each slice. One should select these parameter values intermediate between the extreme values which completely divide the sample set into isolated clusters and those which completely merge the sample set into a single cluster. In practice a narrow range of such values exists.

Applying the Mapper algorithm in this way is an *ad hoc* (case by case) form of ascertaining persistent topological features. *Persistence* is meant in roughly the same sense as the technique of *persistent homology*. Our setting is seemingly inamenable to the application of existing persistent homology tools, both because our setting is multi-dimensional in that the simplicial complexes of interest depend on multiple parameters *n*_*f*_ and *t*, and because a well-defined relation of containment or mapping between the complexes across parameter values is not apparent. Nevertheless persistent features are frequently salient on an *ad hoc* basis, strongly guiding the selection of appropriate values for *n*_*f*_ and *t*.

**Figure 3:**
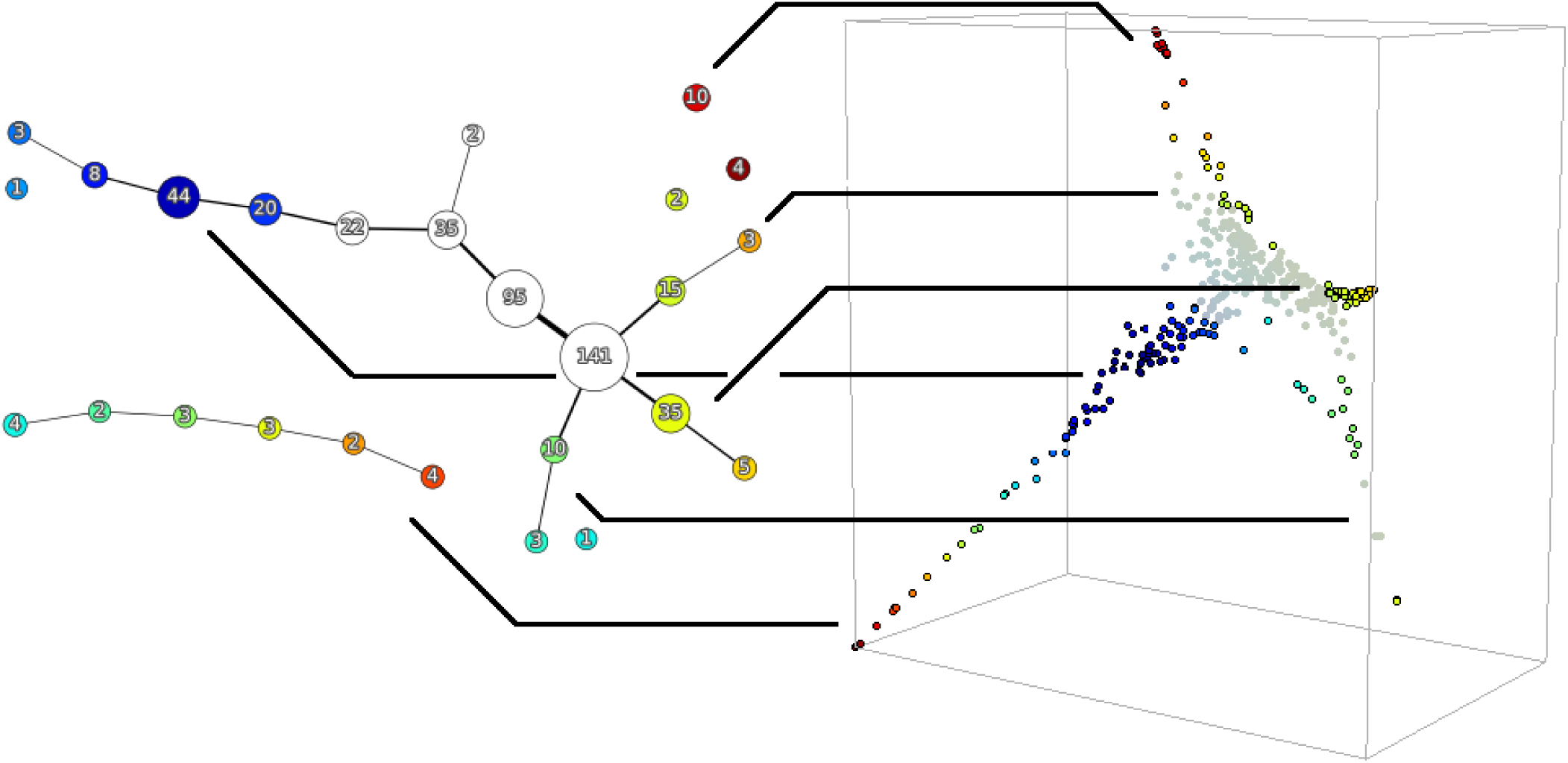
The Mapper state graph of the diffusion re-mapped 355-sample GTEx dataset of Figure 2, with branches highlighted. For illustration, the “core” is not highlighted. The numerical labels indicate the number of samples in a cluster. The color indicates the value of the filter function which was used to seed the Mapper algorithm (a nearest-neighbor network closeness centrality in this case).

### 2.6 State graph extraction and processing

Next, we consider the graph which is the 1-skeleton of the simplicial complex resulting from the Mapper algorithm. We decompose it into linear paths, and concatenate these paths for display.

### 2.7 Plotting heat maps and discerning coherent states

We order the samples within each node of the state graph according to the filter values. This ordering is combined with the concatenated linear path structure for an ordering of the samples *S* along one or both axes of a two-dimensional plot of:

- the expression values
- the correlations with subpopulations defined by discrete covariates
- 1-Wasserstein distance matrix
- diffusion map Euclidean distance matrix

#### 2.7.1 Identify phenotypes

Salient coherent states may appear in the expression heat map defined by patterns of activation and inactivation of particular genes, especially near the two extreme values for the filter function. This may require approximate dichotomization of the expression values (i.e. increasing the contrast, in the terminology of image processing).

From the point of view of topological data analysis, the most interesting states are ones which are not separated by the chosen filter function alone, but are nevertheless distinguished by branching of the state graph. We caution that although the topological aspect of this pipeline has the benefit of insensitivity to dimensionality, functioning well even in very high-dimensional settings, it sometimes provides little insight beyond that already provided by the spectral analysis or diffusion map when the number of samples is small. (On the other hand, Mapper seems to have the potential to function well even without diffusion maps preprocessing or gene network analysis, provided that a filter function can be chosen that brings sufficiently rich outside information into the analysis.)

### 2.8 Interpretation of results

Interpretation of the results or further analysis may include the following:

1. The pattern of activation and inactivation for particular coherent states in the context of the gene network may point to the functioning or non-functioning of components or sub-circuits not previously observed on account of conflation of these states. For example, it could be the case that component *A* is functioning and *B* is non-functioning in state 1, with the opposite behavior in state 2. Considering the whole dataset, before knowing the difference between states 1 and 2, one would only be able to conclude that both *A* and *B* are functioning or partially-functioning.
2. New linkages between components may be hypothesized if sub-circuits *A* and *B*, not apparently directly connected by any process, always function or malfunction together in any given state.
3. The identified states could be used to calibrate the parameters of a dynamic gene expression network model. For example by preferring parameters for which the neighborhoods of these states have high probability of eventual occurrence, or for which these states are stationary.
4. Correlation between patterns of activation/inactivation and additional observations, like drug sensitivity measurements, may help guide or limit the search for causal mechanisms.

## 3 Results

The application of the network-metric/diffusion-map/Mapper pipeline to the 265 gene expression profiles of the samples of the TCGA sarcoma project demonstrates the basic efficacy of the method.

**Figure 4:**
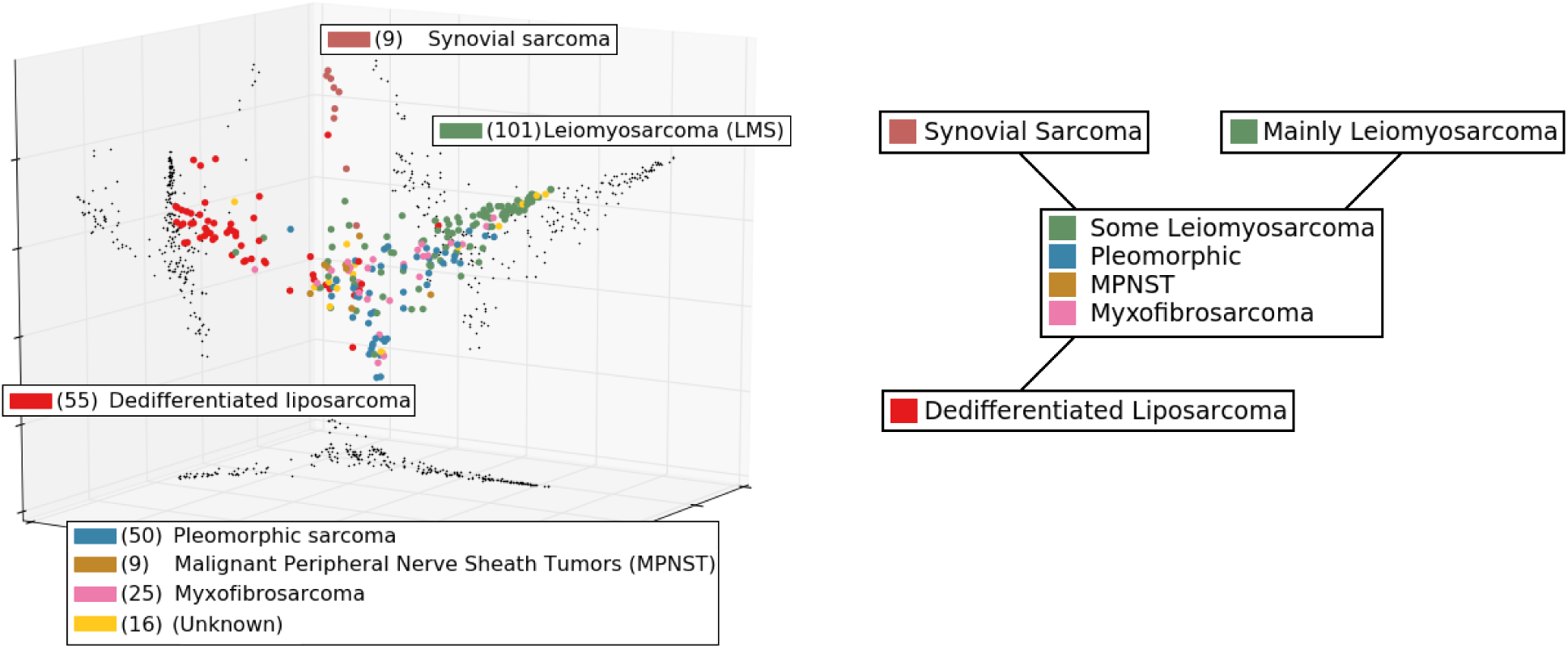
(Left) The diffusion re-mapped images of the 265 gene expression profiles from the TCGA sarcoma project, restricted to the p53 signaling network defined in the KEGG database. The first, second, and fourth eigenvectors were used. (Right) A schematic of the state graph summary produced by the Mapper algorithm.

**Figure 5:**
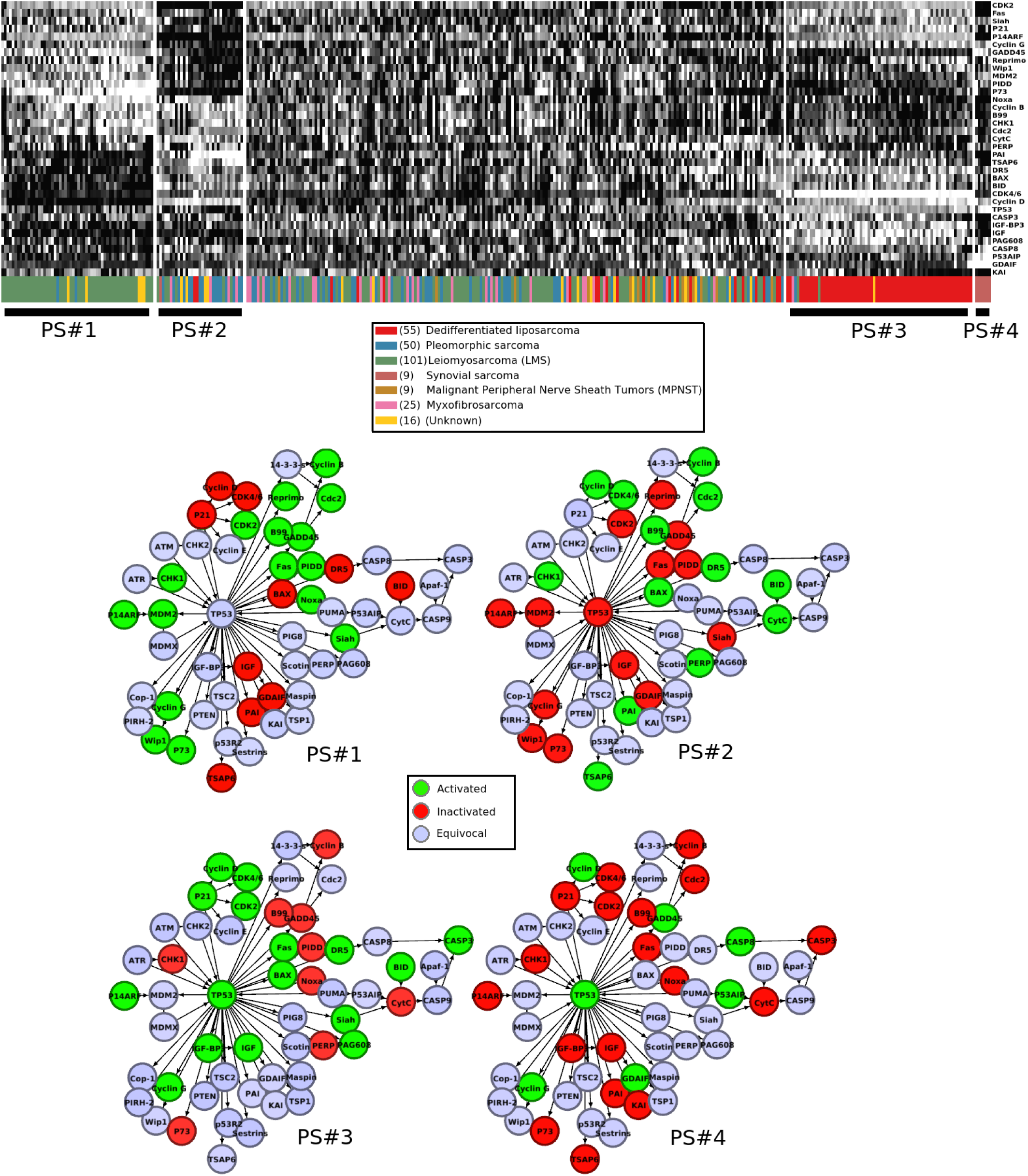
Four coherent states of the KEGG p53 signaling network displayed by subsets of the TCGA sarcoma samples, shown superimposed on the network. See Table 1 for the list of genes.

**Table 1:** The p53 network genes appearing in Figure 5

|  |
| --- |
| CDK2,Fas,Siah,P21,P14ARF,Cyclin G,GADD45,Reprimo,Wip1,MDM2,PIDD,P73<br>Noxa,Cyclin B,B99,CHK1,Cdc2,CytC,PERP,PAI,TSAP6,DR5,BAX,BID,CDK4/6<br>Cyclin D,TP53,CASP3,IGF-BP3,IGF,PAG608,CASP8,P53AIP,GDAIF,KAI |

### 3.1 Discussion of the sarcoma states in Figure 5

We refer to the KEGG p53 signaling pathway at http://www.genome.jp/kegg-bin/show_pathway?map=hsa04115.) The first state PS#1 (P53 Signaling 1) consists almost entirely of samples tagged for leiomyosarcoma, meaning that the corresponding tissue pathology determined a derivation from smooth muscle cells. As expected, the high levels of cyclin G, Wip1, and p73, as well as MDM2 are all negatively regulating TP53, which is not substantially activated. Arrest of the G1 and G2 phases should be triggered since CDK2, cyclin B, and CDC2, are all activated. Although FAS, PIDD, NOXA, and SIAH are substantially activated, the apoptosis pathway for which they are precursors is very much inactivated, including low levels of BAX, the death receptor protein DR5, BID, cytochrome c (CytC), and all caspases. Thus apoptosis is evaded. A plausible hypothesis is that the subpopulation in state PS#1 consists of cells driven to senescence.

The state PS#2 does not consist mainly of any one histopathological subtype. The most obvious feature is that TP53 and almost all of its normal positive regulation targets are inactivated, despite high levels of CHK1 potentially indicating DNA damage. Oddly, nearly all of the markers for TP53 negative feedback regulation are strongly inactivated, including cyclin G, Wip1, p73, and MDM2 seemingly indicating that the feedback system which might normally operate to stabilize levels of TP53 and those of its immediate neighbors is not operating. This is quite the opposite of the situation in state PS#1. Again G1 phase arrest may be triggered, this time because of high levels of cyclin D and CDK4/6 rather than CDK2 (which is substantially inactivated). As expected, P21 loss allows cyclin D and CDK4/6 to appear. With respect to the upstream elements of the apoptosis signaling pathway, we observe in state PS#2 the opposite behavior from the state PS#1, namely that FAS, PIDD, SIAH, and possibly NOXA are lost, but BAX, PERP, and DR5 are activated, as well as BID, which seems to be triggering the release of mitochondrial cytochrome c (CytC). Nevertheless, none of the elements of the caspase cascade are activated and apoptosis still seems to be evaded somehow. Thus state PS#2 reaches a senescence-like outcome substantially similar to that of state PS#1, albeit by apparently very different means.

The third state PS#3 consists almost entirely of samples tagged for dedifferentiated liposarcoma (DDLPS), and conversely almost all of the dedifferentiated liposarcomas among the 265 samples display state PS#3. We emphasize for clarity that all of the states were determined in an entirely unsupervised manner, with no input from the known classification labels. TP53 is strongly activated. Its negative regulator MDM2 seems to be repressed by ARF. A large number of the elements of the normal apoptosis signaling pathway are activated: FAS, BAX, DR5, BID, PAG608, SIAH (without cytochrome c (CytC), however). The very down-stream caspase CASP3 is substantially activated. Apparently, apoptosis does not seem to have been evaded; potentially the dedifferentiated tumors contain the remnants of a large number of cells killed by apoptosis, or, conceivably, necrotic remnants instead; see [Kroemer et al., 2009] for discussion of the possibility of processes blending apoptosis and necrosis. Alternatively, see [Garrido and Kroemer, 2005] for discussion of situations where normally apoptotic caspases are non-lethal to the cell. Substantial activation of P21 some-how isn′t inhibiting CDK4/6 or CDK2; rather CDK4/6 over-expression is the most salient characteristic of state PS#3. According to [Binh et al., 2005], over-expression of CDK4 *and* MDM2 is known to be a reliable diagnostic marker for well-differentiated liposarcoma (not represented in the TCGA sarcoma project).

Note that both leiomyosarcoma and DDLPS subtypes are known to exhibit complex karyotypes, with no apparent characteristic mutation. This seems to be part of the reason why they were selected for inclusion in the TCGA sarcoma project^2^. Nevertheless the coherent states PS#1 and PS#3 show that the expression profiles for these subtypes are more organized than their mutational profiles. We remark that ordinary unsupervised hierarchical clustering substantially reproduces these results, with somewhat less coherence among the apparent states.

On the other hand, synovial sarcoma is known to be well-characterized by a specific translocation resulting in gene fusion of SYT with either SSX1, SSX2, or SSX4 ([Mendelsohn et al., 2015]). So it is not surprising that there is a coherent state, PS#4, displayed by precisely the synovial sarcomas.

## 4 Conclusion and future research directions

We have demonstrated a detailed, unbiased process for identifying clusters of similar disease cases while also identifying similarity via a topological map. One promising direction for future research is the inference of phylogenetic trees via mutational data like Single Nucleotide Polymorphism (SNP) calls or gene amplifications and deletions in the evolutionary context. This context could be spatially dense single-tumor samples or single-patient metastasis or micrometastasis samples. Mapper would be especially adapted to the elucidation of branching/inheritance structures when the filter function is a suitable quantification of the deviation of a sample from a founder population. For example, a Hamming-type distance in the case of SNP sequences, which could also be used for the intrinsic metric between SNP sequences. As an alternative to the naive Hamming distance, a network-enriched distance could be obtained by means of linkage disequilibrium calculations.

## 5 Acknowledgements

We thank Daniel Müllner for making his Python Mapper implementation so readily available (http://danifold.net/mapper/). We also thank Juan M. Bello-Rivas for the use of his diffusion maps software (https://github.com/jmbr/diffusion-maps). C.M. thanks the Department of Chemical and Biological Engineering, Princeton University, for their hospitality. This project was supported by AFOSR grant FA9550-17-1-0435), RO grant (W911NF-17-1-049), grants from National Institutes of Health (1U24CA18092401A1, R01-AG048769), MSK Cancer Center Support Grant/Core Grant (P30 CA008748), and a grant from Breast Cancer Research Foundation (grant BCRF-17-193). IGK and CM thank the National Science Foundation for partial support.

## Footnotes

1 http://danifold.net/mapper/

2 https://cancergenome.nih.gov/cancersselected/Sarcoma

## References

Binh, M. B., Sastre-Garau, X., Guillou, L., de Pinieux, G., Terrier, P., Lagace, R., Aurias, A., Hostein, I., and Coindre, J. M. (2005). MDM2 and CDK4 immunostainings are useful adjuncts in diagnosing well-differentiated and dedifferentiated liposarcoma subtypes: a comparative analysis of 559 soft tissue neoplasms with genetic data. Am. J. Surg. Pathol., 29(10):1340–1347.

Camara, P. G., Rosenbloom, D. I., Emmett, K. J., Levine, A. J., and Rabadan, R. (2016). Topological data analysis generates high-resolution, genome-wide maps of human recombination. Cell Syst, 3(1):83–94.

Chen, Y., Cruz, F. D., Sandhu, R., Kung, A. L., Mundi, P., Deasy, J. O., and Tannenbaum, A. (2017). Pediatric sarcoma data forms a unique cluster measured via the Earth Mover’s Distance. Sci Rep, 7(1):7035.

Coifman, R. R. and Lafon, S. (2006). Diffusion maps. Applied and Computational Harmonic Analysis, 21(1):5–30. Special Issue: Diffusion Maps and Wavelets.

Coifman, R. R., Lafon, S., Lee, A. B., Maggioni, M., Nadler, B., Warner, F., and Zucker, S. W. (2005). Geometric diffusions as a tool for harmonic analysis and structure definition of data: Diffusion maps. PNAS, 102(21):7426–7431.

Edelsbrunner, H. and Harer, J. L. (2010). Computational Topology. American Mathematical Society, Providence, RI.

Garrido, C. and Kroemer, G. (2005). Life’s smile, death’s grin: Vital functions of apoptosis-executing proteins. Current opinion in cell biology, 16:639–46.

James, R. G., Davidson, K. C., Bosch, K. A., Biechele, T. L., Robin, N. C., Taylor, R. J., Major, M. B., Camp, N. D., Fowler, K., Martins, T. J., and Moon, R. T. (2012). WIKI4, a novel inhibitor of tankyrase and Wnt/-catenin signaling. PLoS ONE, 7(12):e50457.

Jones, P. W., Maggioni, M., and Schul, R. (2008). Manifold parametrizations by eigenfunctions of the laplacian and heat kernels. Proceedings of the National Academy of Sciences, 105(6):1803–1808.

Kroemer, G., Galluzzi, L., Vandenabeele, P., Abrams, J., Alnemri, E. S., Baehrecke, E. H., Blagosklonny, M. V., El-Deiry, W. S., Golstein, P., Green, D. R., Hengartner, M., Knight, R. A., Kumar, S., Lipton, S. A., Malorni, W., Nuez, G., Peter, M. E., Tschopp, J., Yuan, J., Piacentini, M., Zhivotovsky, B., and Melino, G. (2009). Classification of cell death: recommendations of the nomenclature committee on cell death. Cell Death And Differentiation, 16(3).

Lockwood, S. and Krishnamoorthy, B. (2014). Topological features in cancer gene expression data. http://arxiv.org/abs/1410.3198v1.

Mendelsohn, J., Howley, P. M., Israel, M. A., Gray, J. W., and Thompson, C. B. (2015). Molecular Basis of Cancer. Elsevier, 4 edition.

Munkres, J. R. (1975). Topology: A First Course. Prentice-Hall, Inc., Englewood Cliffs, N.J.

Nadler, B., Lafon, S., Coifman, R. R., and Kevrekidis, I. G. (2006). Diffusion maps, spectral clustering and reaction coordinates of dynamical systems. Applied and Computational Harmonic Analysis, 21(1):113–127. Special Issue: Diffusion Maps and Wavelets.

Nicolau, M., Levine, A. J., and Carlsson, G. (2011). Topology based data analysis identifies a subgroup of breast cancers with a unique mutational profile and excellent survival. Proc. Natl. Acad. Sci. U.S.A., 108(17):7265–7270.

Rachev, S. T. and Rüschendorf, L. (1998). Mass transportation problems. Vol. II. Probability and its Applications (New York). Springer-Verlag, New York. Applications.

Rajendran, K., Kattis, A., Holiday, A., Kondor, R., and Kevrekidis, I. G. (2016). Data mining when each data point is a network. https://arxiv.org/abs/1612.02908.

Rubner, Y., Tomasi, C., and Guibas, L. J. (2000). The earth mover’s distance as a metric for image retrieval. International Journal of Computer Vision, 40(2):99–121.

Seemann, L., Shulman, J., and Gunaratne, G. H. (2012). A robust topology-based algorithm for gene expression profiling. ISRN Bioinform, 2012:381023.

Xu, R., Damelin, S., Nadler, B., and Wunsch, D. C. (2010). Clustering of high-dimensional gene expression data with feature filtering methods and diffusion maps. Artificial Intelligence in Medicine, 48(2):91–98. Artificial Intelligence in Biomedical Engineering and Informatics.

